# New gene association measures by joint network embedding of multiple gene expression datasets

**DOI:** 10.1101/2020.03.16.992396

**Authors:** Guiying Wu, Xiangyu Li, Wenbo Guo, Zheng Wei, Tao Hu, Jin Gu

**Author notes:** To whom correspondence should be addressed. (Ext. 866).

## Abstract

Large number of samples are required to construct a reliable gene co-expression network, the samples from a single gene expression dataset are obviously not enough. However, batch effect may widely exist among datasets due to different experimental conditions. We proposed JEBIN (Joint Embedding of multiple BIpartite Networks) algorithm, it can learn a low-dimensional representation vector for each gene by integrating multiple bipartite networks, and each network corresponds to one dataset. JEBIN owns many inherent advantages, such as it is a nonlinear, global model, has linear time complexity with the number of genes, dataset or samples, and can integrate datasets with different distribution. We verified the effectiveness and scalability of JEBIN through a series of simulation experiments, and proved better performance on real biological data than commonly used integration algorithms. In addition, we conducted a differential co-expression analysis of hepatocellular carcinoma between the single-cell and bulk RNA-seq data, and also a contrast between the hepatocellular carcinoma and its adjacency samples using the bulk RNA-seq data. Analysis results prove that JEBIN can obtain comprehensive and stable gene co-expression networks through integrating multiple datasets and has wide prospect in the functional annotation of unknown genes and the regulatory mechanism inference of target genes.

## INTRODUCTION

Genes involved in normal and diseased life process generally do not function alone, but rather determine the biological phenotype through regulatory or collaborative interactions among a group of genes [1,2]. The functionally related genes tend to exhibit co-expression relationships [3-5] in gene expression data, named “guilt-by-association” (GBA) [6]. Gene co-expression networks are powerful tools for describing the overall interactions among genes and imply the causal information which can be used to infer regulatory relationships [7,8]. The annotation information about the unannotated genes, such as the biological pathways they may participate in and the functions they possibly possess, can be obtained based on the connections between the unannotated genes and the well investigated genes in their co-expression network neighbourhood [9-11]. Since thousands of biological samples are required to construct a trustworthy gene co-expression network [12,13], the samples from a single gene expression dataset are obviously not enough [14], it becomes an urgent problem to integrate multiple datasets so as to build a reliable gene co-expression network. Megan Crow et al. demonstrated that for both mouse bulk and single-cell RNA-seq data, the co-expression networks obtained by integrating multiple datasets performed better than that obtained by using only a single dataset [15].

Traditional gene association measure methods are designed to calculate the similarity between the expression vectors of two genes directly, and the commonly used methods include Pearson correlation coefficient, Spearman correlation coefficient, Kendall correlation coefficient, biweight midcorrelation and mutual information, etc. [16-18]. Since the gene sets corresponding to different datasets may be inconsistent, the integration of multiple gene adjacent matrices for each dataset inevitably produces missing values by calculating the minimum [19], average, weighted average [20] or other statistic values for each corresponding matrix element. A large amount of missing values can affect the effect of matrix imputation conducted before the integration of gene adjacent matrices [21]. It should be noted that when constructing a gene co-expression network, the above methods calculate the correlation between the current two genes, without taking into account information from other genes [16]. WGCNA (Weighted gene co-expression network analysis) [19,22] is a commonly used method designed primarily to achieve the gene modules. WGCNA also provides the one-step method for the co-expression network integration of multiple datasets, but the identical gene sets from different datasets are the necessary condition for the input. GeneMANIA is another commonly used method to integrate multiple gene association networks [20]. The ridge regression algorithm is implemented to calculate the weights of different association networks, then the weighted average of these networks is carried out to get the integrated network, on which the function of genes can be predicted by the label propagation algorithm. This algorithm also requires the the same set of genes across different association networks by default. Moreover, in order to keep the sparsity of the networks, only the top 50 gene association values were reserved for each gene, and the other values were assigned to 0. To avoid the problem of different gene sets in different datasets, there are also studies conducted the integration operations at the level of gene pairs rather than the adjacency matrix [23]. After calculating the correlation value among all gene pairs in each dataset, only the pairs appearing in at least N datasets were selected for the integration to construct the final gene association network. The disadvantage of these methods is that some genes may miss important correlations and affect subsequent functional analysis, because they can only obtain their association with some genes rather than with all other genes.

The above-mentioned methods calculate the association degree between one gene and another directly to construct the association network for each dataset. However, in real organisms, a gene is likely to interact with a group of genes or even a group of genes interact with another group of genes, thereby making the gene to be studied may not have strong marginal association with the individual gene in a group of genes [16]. Since high-order association may be missed in such paired similarity measurements, another class of algorithms that utilize the Gaussian Graphical models (GGM) attempt to describe the complex gene association networks by the concept of conditional dependence [16,21] [24-28]. GGM is designed to obtain the precision matrix, also known as the inverse covariance matrix or concentration matrix, by maximizing the log likelihood of multivariate Gaussian distribution and controlling the sparseness of the edges by penalizing the sum of the absolute values of the off-diagonal elements in the precision matrix (*L*1 norm), and the non-zero mode in the precision matrix corresponds to a graph structure, reflecting the association of all gene pairs [26,29,30]. There are two levels of application for GGM in the field of gene association analysis, one is the graphical lasso algorithm at the gene level [24], the other is the module graphical lasso algorithm at the level of gene module [21,25]. The graph lasso algorithm assumes the independence among the matrix elements when applying sparsity penalty to the precision matrix, that is the independence of the edges in gene co-expression networks. Nevertheless, the edges are not independent in real gene association networks. Thus, Celik et al. proposed the Modular Graphical Lasso algorithm (MGL) [25], in which correlated genes are considers as a module and the conditional dependence between two modules are modelled as a GGM. However, MGL loses the associations at gene level and is designed for a single dataset. In the field of integrating multiple datasets to construct gene co-expression network, the algorithms with GGM as the main thought include JGL (Joint Graphical Lasso) [24] and TDJGL (Two Dimensional Joint Graphical Lasso) [27] at the gene level, and INSPIRE (INferring Shared modules from multiple gene expression datasets) at the module level [21]. To get the inverse covariance matrix, this class of GGM based methods have merely utilized the linear relationships of gene pairs, because the input empirical covariance matrix measures the linear association of genes only.

Another group of integration methods on multiple gene expression datasets include Generalized Singular Value Decomposition (GSVD) algorithm for two datasets [31] and High-Order Generalized Singular Value Decomposition algorithm (HO GSVD) for multiple datasets [32,33]. The inputs of these algorithms can be either the original expression matrixes or the gene co-expression matrixes. The disadvantage is that only a collection of genes shared by all datasets and the gene sets specific to one dataset are obtained in the results, so the gene level co-expression networks have been missed as well. In addition, they also require that the set of genes corresponding to each dataset must be identical, and the calculation time increases exponentially as the number of genes or the number of datasets increases.

We propose a new model of Joint Embedding of multiple BIpartite Networks (JEBIN) to learn the low-dimensional vector representation of genes utilizing the nonlinear relationships among genes from multiple gene expression data sets. With the genes’ representation vectors learned, gene co-expression networks and gene modules can be obtained using the commonly used vector similarity calculating method and traditional clustering methods separately.

## MATERIAL AND METHODS

### Overview

JEBIN is designed to learn the low-dimensional representation, named the “consensus vector” of genes, by integrating multiple bipartite networks with each corresponding to one gene expression dataset. As shown in Figure 1, the main procedure of JEBIN is to transform each gene expression dataset to a bipartite network, and then joint embedding multiple bipartite network to learn the dataset-specific representation vectors for the genes in each network and the consensus representation vectors for all the genes appeared in at least one dataset. Utilizing the low-dimensional consensus vectors, we can visualize the genes on a 2D view. The gene association networks and gene clustering can also be calculated rapidly.

**Figure 1.**
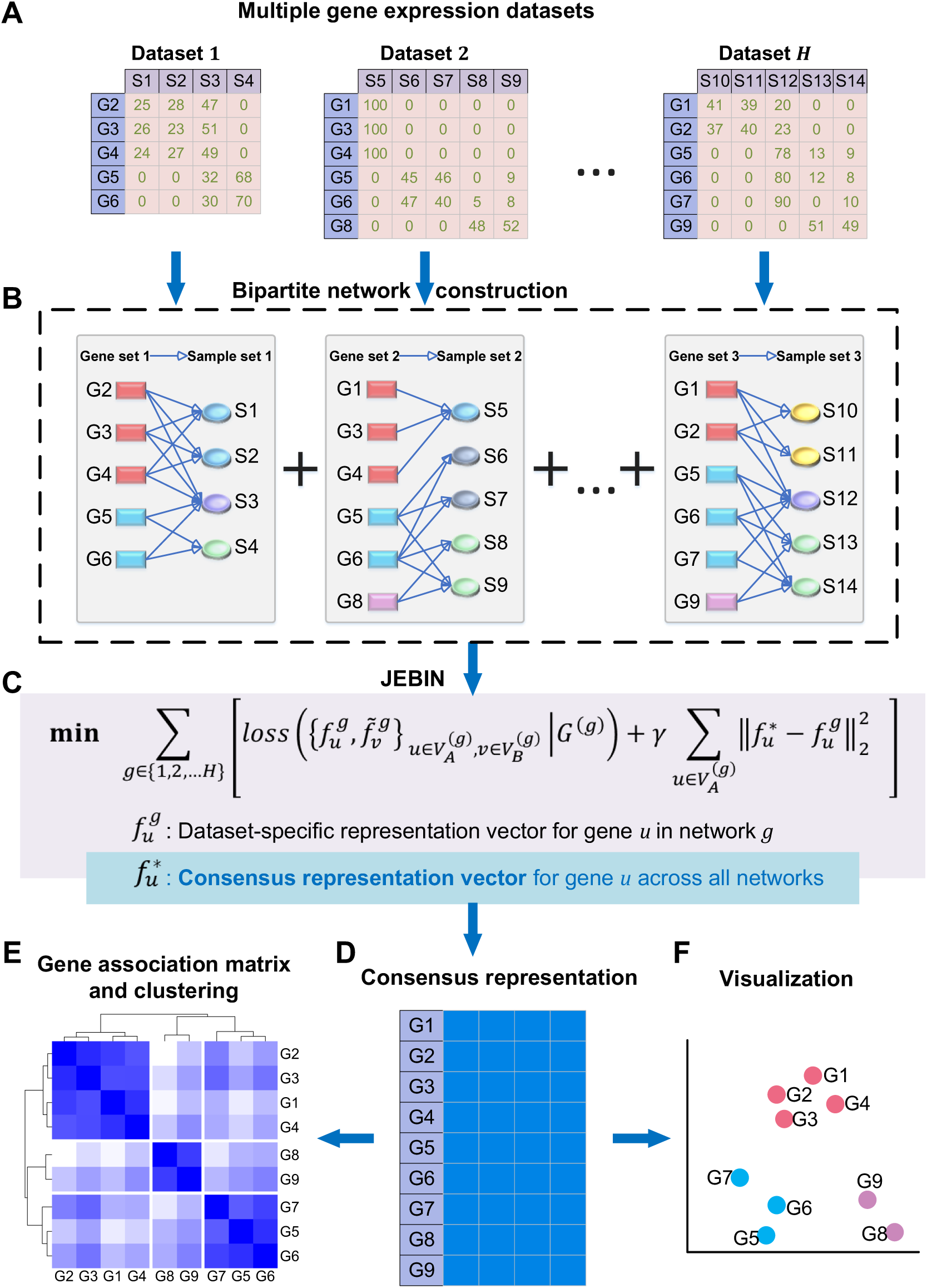
Overview of the pipeline of JEBIN. (A) Input multiple gene expression datasets. (B) Bipartite network construction. (C) The objective function of JEBIN. (D) Consensus representation of genes. (E) Gene association networks construction and gene clustering. (F) Visualization of genes.

### JEBIN model

Given *H* gene expression datasets in matrix form, we can construct a bipartite network graph 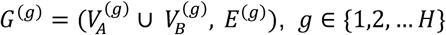, for each dataset [34,35]. 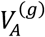 and 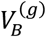 are twodisjoint sets of vertices with different types, the gene set and sample set respectively. *E*^(*g*)^ is the set of edges between 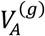 and 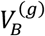, the weight for each edge represents the expression value of the corresponding gene in the linked sample. In our JEBIN model, the gene vertex sets 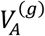 are not required to be completely the same across all bipartite networks, while the sample vertex sets 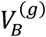 are regarded as totally different sets, with no intersection for different bipartite networks.

We denote 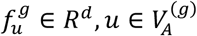 and 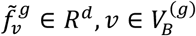 as the identity and context embedding vector, respectively, in network graph *G*^(*g*)^, *g* ∈ {1,2, … *H*}. *d* are the dimensions of the embedding vectors, they are required to be the same in different networks. The object function of joint embedding of all bipartite networks *G*^(*g*)^, *g* ∈ {1,2, … *H*}, is

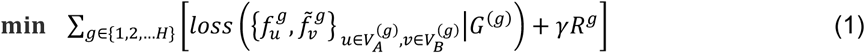

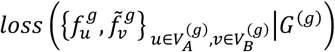 is the intra-graph loss function of network graph *G*^(*g*)^, reflecting the ability of 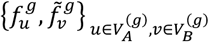 vectors to depict network *G*^(*g*)^, and the second term *γR*^*g*^ regularizes the embedding vectors across different networks.

For each directed edge (*u, v*) ∈ *E*^(*g*)^ in network *G*^(*g*)^, the conditional distribution is defined as the following softmax function:

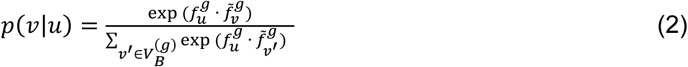

and the empirical distribution is defined as

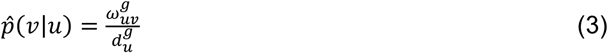

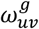 is the weight of the edge (*u, v*) in network *G*^(*g*)^, and 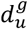 is the out-degree of vertex *u*, calculated by 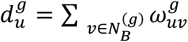, where 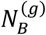is the set of out-neighbours of vertex *u*, which is a subset of 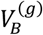 in network graph *G*^(*g*)^.

The weighted summation of distances between the conditional distribution *p*(·|*u*) and the empirical distribution 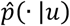 for each node 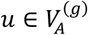 is used to represent the intra-graph loss function:

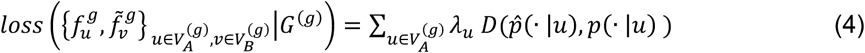

We adopt KL-divergence as the distance function between two distributions, and simply set *λ*_*u*_ as the out-degree of vertex *u, λ*_*u*_ = *d*_*u*_, to reflect the importance of vertex *u* in the network. Omitting some constants, the objective function in equation (4) can be calculated as

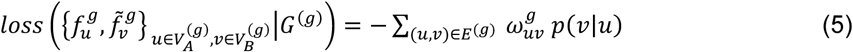

Since *p*(*v*|*u*) is computationally expensive, which requires the summation over the entire set of vertices in 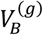 as in equation (2), we adopt negative sampling strategy to address this problem [36]. The main idea of negative sampling is to distinguish the target node *v* from the nodes 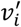 drawn from the noise distribution *P*^(*g*)^(*v*) by logistic regression. The objective function in equation (5) can be rewritten as

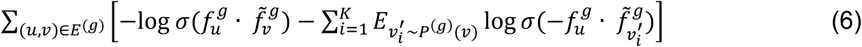

where *σ*(*x*) = 1/(1 + exp (−*x*)) is the sigmoid function, and *K* is the number of negative edges (default 5). The first term models the network edges (*u*^*g*^, *v*^*g*^) ∈ *E*^(*g*)^ sampled from the observed edges, and the second term models the negative edges 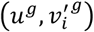 drawn from the noise distribution 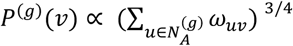, where 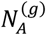 is the set of in-neighbours of vertex *v*.

For joint embedding of multiple bipartite networks, we defined a consensus embedding vector 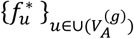 for each vertex *u* in the union set of 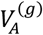, which models the shared information implied in all datasets because of some shared latent factors. The second term *γR*^*g*^ in equation (1) is used to regularize the consensus vectors 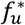 utilizing the dataset-specific vectors 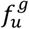, which defined as follows:

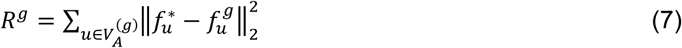

Through this term, we can get the consensus vector for each gene, representing the consensus information from multiple gene expression datasets. The positions of genes in the embedding space can reflect the similarity among the expression profiles of genes.

With negative sampling, the objective function in equation (1) involving one real network edge (*u*^*g*^, *v*^*g*^) ∈ *E*^(*g*)^ and *K* negative edges 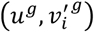, *i* ∈ {1,2, …*K*} drawn from only one network graph *g* is as follows:

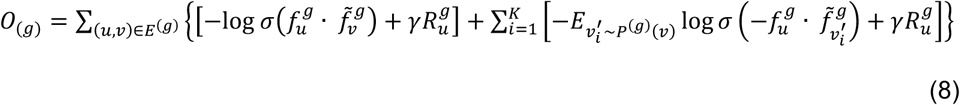

where

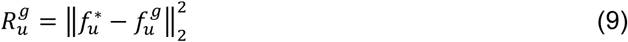

Finally, the objective function in equation (1) equals to the summation of *O*_(*g*)_ across all datasets, written as

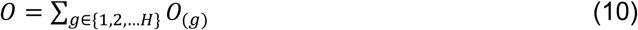

Asynchronous stochastic gradient descent (ASGD) algorithm [37] is adopted to optimize the equation (8). Since the gradient will be multiplied by the weight of the edge, there would be a problem to select a suitable learning rate when the weights of edges have a high variance. Tang have proposed “Edge Sampling” strategy to treat this problem by unfolding an edge with weight *ω* into *ω* binary edges [38]. Alias table method [39] can effectively decrease the time of sampling an edge from *O*(|*E*|) to *O*(|1|). In practice, we set the parameter of edge sampling number to be proportional to the edge number of the bipartite network. Then for each bipartite network, the negative sampling optimization takes *O*(*d*(*K* + 1)|*E*|) time, which is linear to the number of edges |*E*|. For the joint embedding of *H* datasets, the overall time complexity is *O*(*H*|*E*|), here |*E*| represents the maximum edge number of *H* bipartite networks. The edge number of a bipartite network |*E*| is the number of nonzero elements in a gene expression matrix, so our algorithm can be well-suited to sparse matrix data, for instance, single cell RNA-seq data with enormous number of cells and very sparse gene expression values.

JEBIN has four outstanding features. Firstly, the information from the whole gene population is employed to get the consensus representation of each genes across multiple gene expression datasets. That is, the relative position of one gene in the low-dimensional space is affected by all the other genes which makes JEBIN a global method. Secondly, JEBIN can effectively dispose the problem of inconsistent gene sets among different datasets due to different sequencing techniques or pre-processing. The final consensus gene co-expression networks will cover all the genes appeared in at least one dataset, and the consistent and stable co-expression relationships occurring across multiple datasets can be further enhanced. Thirdly, the calculation time of JEBIN increases linearly with the number of genes and also the number of datasets. Fourthly, JEBIN can integrate biological data that is subject to different distributions, as well as has promising application prospects in integrating multi-omics data and fusing biological prior knowledge.

## RESULTS

To verify the effectiveness and scalability of JEBIN, we designed a series of simulation experiments, and conducted the comparison of JEBIN with commonly used integration algorithms on real biological data.

### Simulation experiments

We conducted simulation experiments with small-scale gene networks designed as the co-expression networks as in Figure 2A and the gene regulatory network as in Figure 2B. And then Gaussian Graphical models and thermodynamic model is used to generate the gene expression data which contain linear and nonlinear relationships among genes respectively.

**Figure 2.**
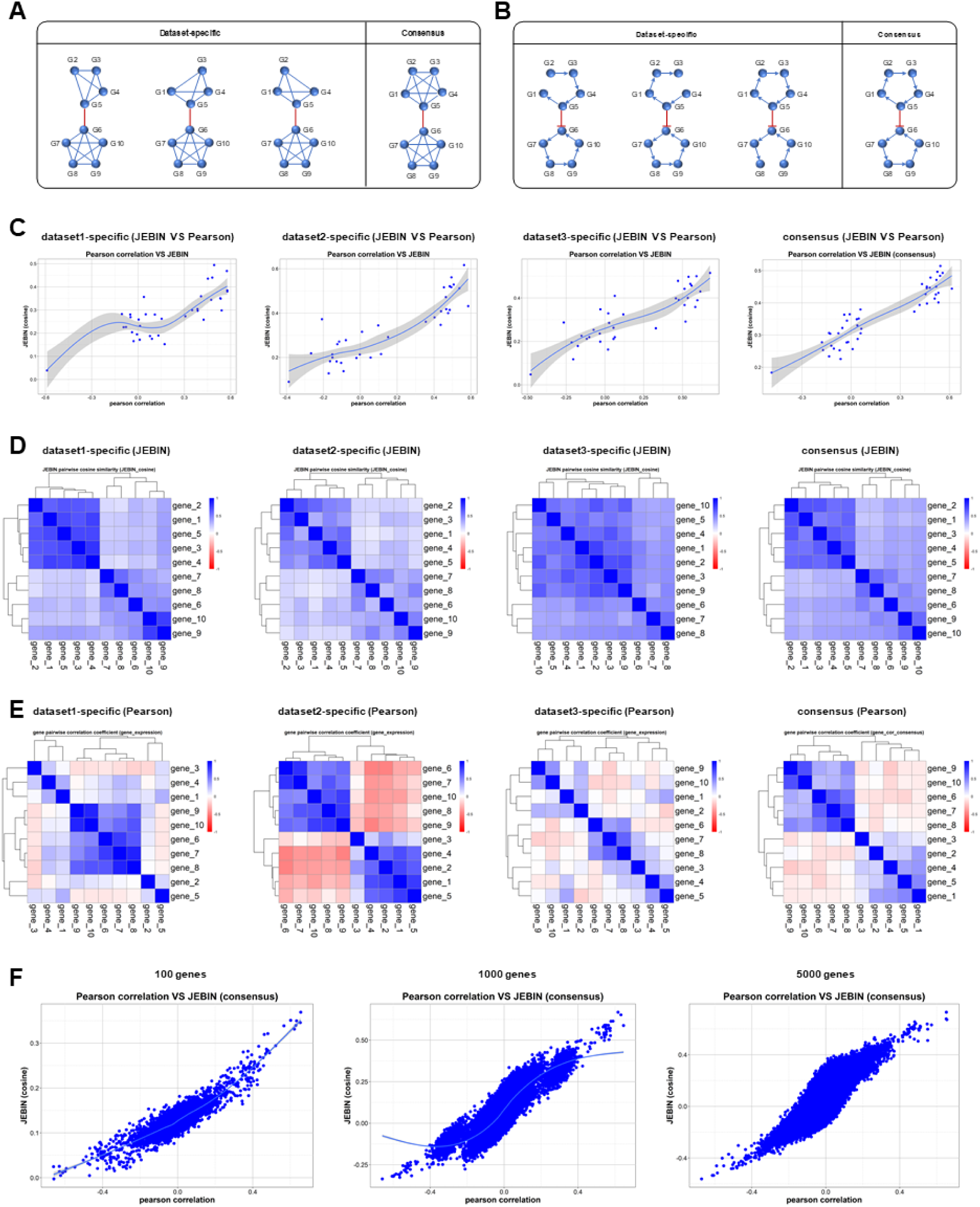
Simulation experiments. (A) Co-expression networks and (B) gene regulatory network designed to generate simulative gene expression data. (C) Compare the cosine similarity between the vector representations of gene pairs obtained by JEBIN with Pearson correlation integration methods. (D) The cosine similarity between the vector representations of gene pairs obtained by JEBIN and genes clustering. (E) The Pearson correlation integrated similarity and genes clustering. (F) The cosine similarity between the gene pairs obtained by JEBIN consistently maintained a good linear correlation with Pearson correlation integration methods.

For the linear simulation data, the covariance matrix among genes was set, and then the observation samples were generated by using the multivariate gaussian distribution. So, the Pearson correlation coefficients between gene pairs can be used as the benchmark measurement. The cosine similarity between the vector representations of gene pairs obtained by JEBIN is calculated, and the results had shown good linear relationship with the Pearson correlation coefficients, as shown in Figure 2C. This demonstrates that JEBIN can efficiently uncover the linear positive co-expression relationships in small-scale gene networks.

For the nonlinear simulation data, the thermodynamic model was used to generate the observation samples based on the gene regulatory network shown in Figure 2B. The cosine similarity between the vector representations of gene pairs obtained by JEBIN was calculated and clustering was conducted. It was found that the clustering results as shown in Figure 2D, the consensus gene representation vectors could correctly identify two gene modules with positive internal correlation. Furthermore, for dataset-specific clustering, the results of JEBIN are more stable than the results of Pearson correlation which is shown in Figure 2E. This indicates that JEBIN can identify the nonlinear correlation between gene pairs.

Then we expanded to large-scale gene networks. The linear relationships are still modelled by Gaussian Graphical models. But for the nonlinear relationships, we used multiple real gene expression datasets.

When using the gaussian graph model, the number of integrated datasets was expanded to 10, and the number of genes was set to 100, 1000 and 5,000 genes, respectively. The results shown in Figure 2F indicated that the cosine similarity between the gene pairs obtained by JEBIN consistently maintained a good linear correlation with Pearson correlation coefficient, which verified the effectiveness of JEBIN on linear relationships mining in large-scale gene networks.

### Real biology data

We collected the datasets from six groups of different conditions, including hepatocellular carcinoma (HCC), the adjacent data of HCC, breast cancer, ovarian cancer, colorectal cancer and bladder cancer. JEBIN was used to integrate multiple datasets from different experiments, and the learned consensus vectors of genes was visualized by tSNE, as shown in Figure 3A. The aggregation of genes can be clearly seen, suggesting that these representation vectors can be used for the discovery of gene modules. We compared JEBIN with four commonly used methods for calculating genetic correlations, Pearson correlation, Spearman correlation, biweight midcorraltion, and cosine similarity, and found that JEBIN consistently performed best in all 6 experiments, as is shown in Figure 3B.

**Figure 3.**
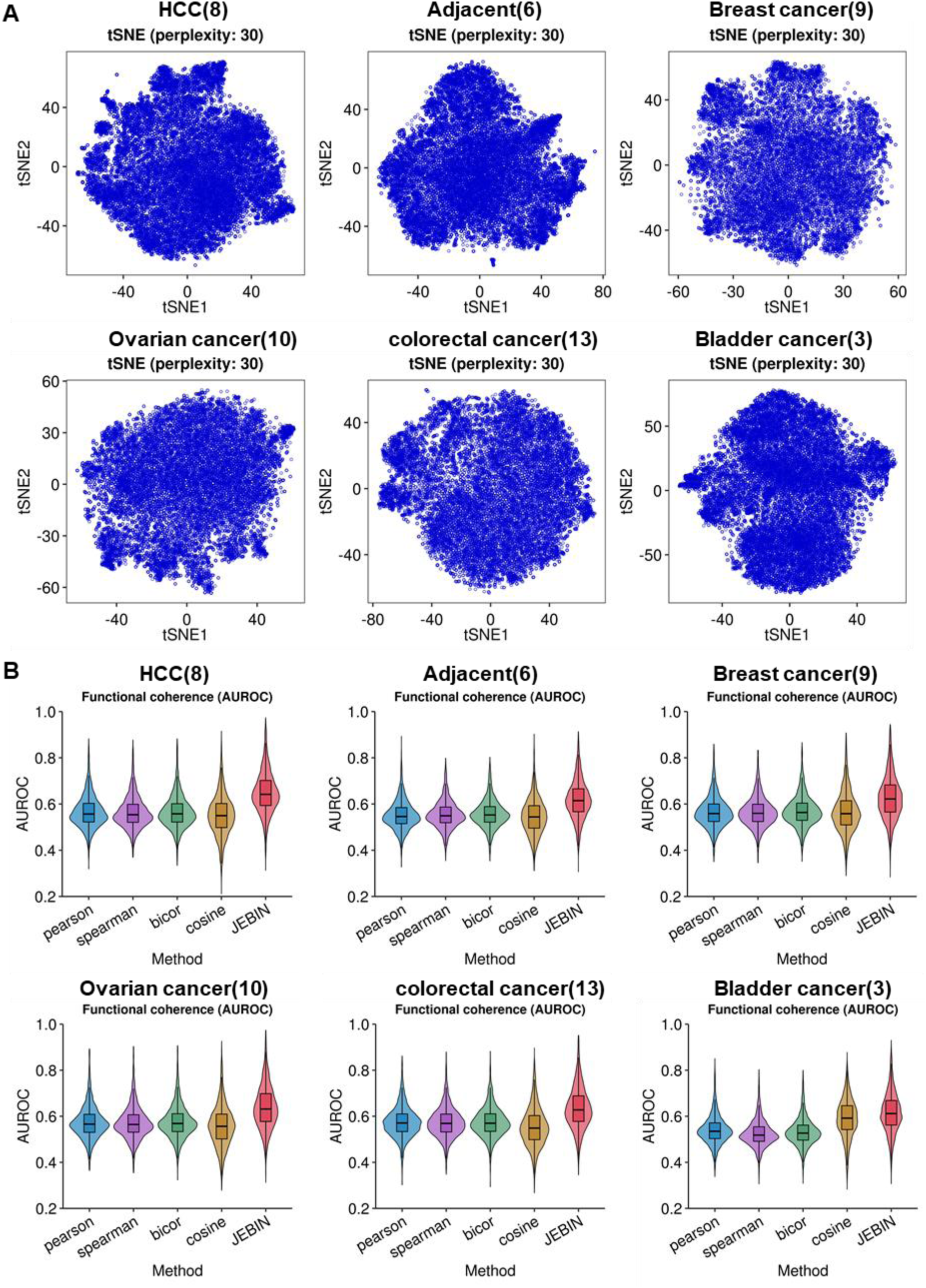
Visualization of genes calculated by JEBIN on real biology data (3A) and performance contrast with four commonly used methods (3B).

### Differential co-expression analysis

In the follow-up application, we conducted a differential co-expression analysis of hepatocellular carcinoma between the single-cell and bulk RNA-seq data, and also a contrast between the hepatocellular carcinoma and its adjacency samples using the bulk RNA-seq data. Results shown in Figure 4A is the enrichment results of HCC-specifc co-expression genes when compared with the adjacent bulk RNA-seq data, the top is the enrichment on GO BP terms, and the bottom is the enrichment on KEGG pathways. DNA repair and DNA replication terms are all found in both enrichments, which is consistent with our knowledge that abnormalities often occur in cancers. This result also proves the effectiveness of JEBIN on the association calculation of genes.

**Figure 4.**
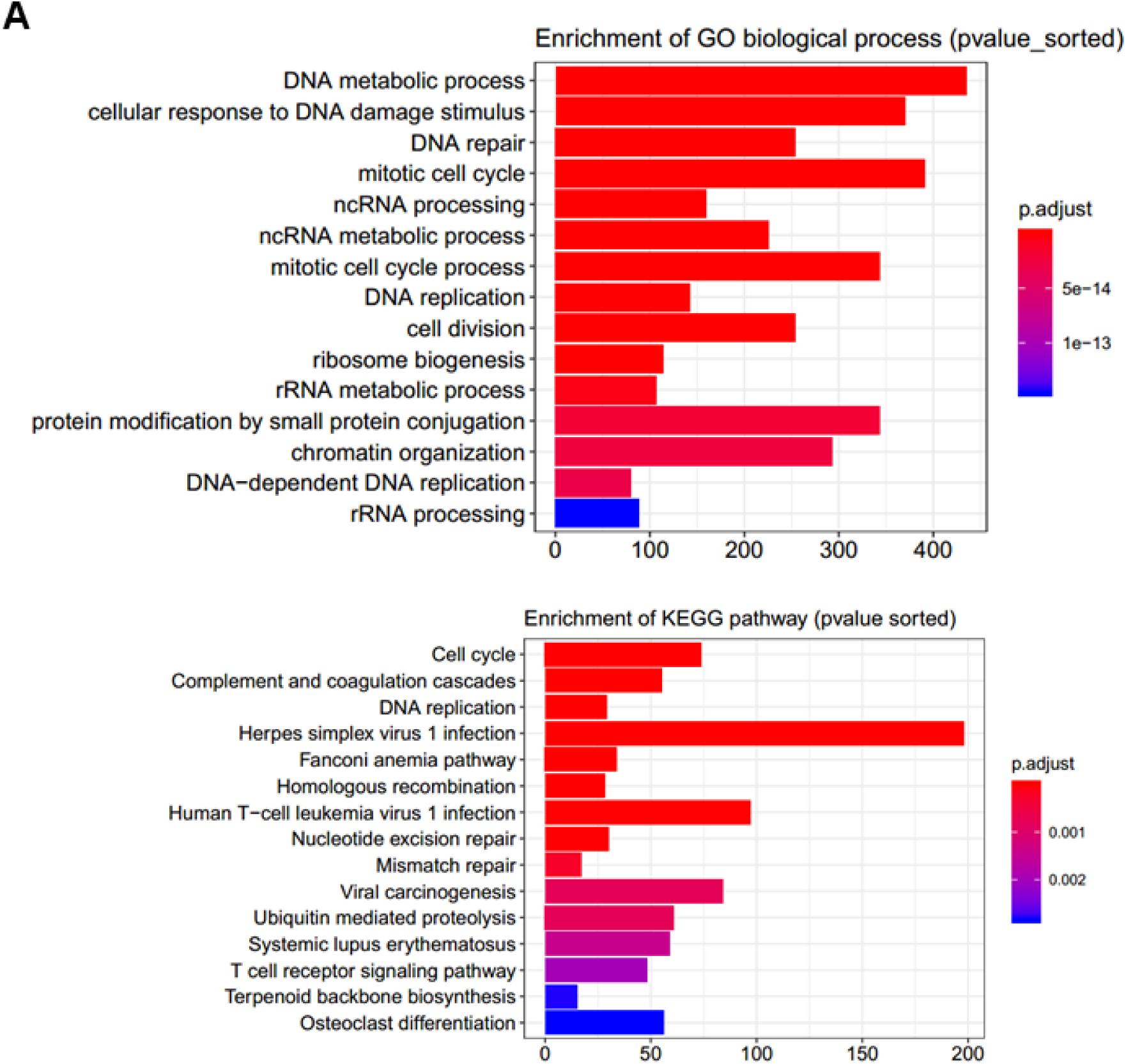
Enrichment results of HCC-specifc co-expression genes when compared with the adjacent bulk RNA-seq data. The top is the enrichment on GO BP terms, and the bottom is the enrichment on KEGG pathways.

## DISCUSSION

Multiple gene expression datasets contain consensus information because of common latent factors, and dataset-specific information for condition-specific factors. Furthermore, different datasets can provide complementary information, so the integration of them can generate more complete and stable gene co-expression networks. Our proposed method, JEBIN, can construct the consensus gene co-expression network by joint embedding multiple datasets from different platforms or studies. JEBIN owns many inherent model advantages: (1) nonlinear model, it can effectively dig the linear and nonlinear correlation between gene pairs, (2) global algorithm, it utilizes the expression data of all genes to determine the representation vector of each gene, (3) the time complexity is linearly related to the number of genes, samples and datasets respectively, (4) JEBIN can integrate datasets of different distributions, such as microarray data and RNA-seq data. The effectiveness and scalability of JEBIN were verified through a series of simulation experiments, and showed better performance on real biological data than commonly used integration algorithms. The differential co-expression analysis is conducted on hepatocellular carcinoma between the single-cell and bulk RNA-seq data, and on the bulk RNA-seq data between the hepatocellular carcinoma and its adjacency samples. Analysis results prove that JEBIN has good prospect in the functional annotation of unknown genes and the regulatory mechanism inference of target genes.

JEBIN belongs to the class of representation learning methods. Representation learning is a rapidly developed class of algorithms which can effectively deal with large-scale networks through integrating the information from multiple sources and learning the vector representation for each studied object. In the future, it will be a powerful tool for the integration of various level of biological data, like muti-omics data. And its application in the differential co-expression analysis between the disease and the normal datasets can provide a possible target for the treatment of diseases.

## AUTHOR’S CONTRIBUTIONS

G.Y.W. and J.G. initiated the project. G.Y.W. developed the method and complete the data analysis. G.Y.W., X.Y.L. and Z.W. wrote the codes. W.B.G. and T.H. did some data preprocessing. G.Y.W., X.Y.L., and J.G. wrote the manuscript. All authors read and approved the final manuscript.

